# Capsular polysaccharide-mediated protein loading onto extracellular membrane vesicles of a fish intestinal bacterium, *Shewanella vesiculosa* HM13

**DOI:** 10.1101/2023.04.25.538355

**Authors:** Kouhei Kamasaka, Jun Kawamoto, Taiku Tsudzuki, Yuying Liu, Tomoya Imai, Takuya Ogawa, Tatsuo Kurihara

## Abstract

Bacterial extracellular membrane vesicles (EMVs) play various physiologically important roles mediated by cargo proteins. However, our understanding of the molecular mechanism underlying cargo loading onto EMVs is limited. In this study, we analyzed the mechanism of cargo protein loading onto EMVs from a fish intestinal Gram-negative bacterium, *Shewanella vesiculosa* HM13. This strain secretes EMVs carrying a major cargo protein, P49. Near the P49 gene, we found genes having homology to genes involved in protein secretion and surface polysaccharide-chain synthesis. Among them, the deletion of genes encoding homologs of a flippase involved in bacterial extracellular polysaccharide synthesis (HM3343), phosphoethanolamine transferase (HM3344), and glycerophosphodiester phosphodiesterase (HM3345) resulted in the loss of capsular polysaccharide (CPS) of EMVs. We conducted an *in vitro* P49 loading assay onto P49-free EMVs to examine whether P49 was loaded onto the EMVs via its interaction with the CPS of the EMVs. We found that purified P49 was loaded onto EMVs harboring CPS *in vitro*, whereas it was not loaded onto EMVs from the mutants lacking CPS production due to the loss of HM3343, HM3344, and HM3345. Transmission electron microscopy of EMVs loaded with P49 *in vitro* and *in vivo* showed spherical nanoparticles around the EMVs, whereas such particles were not observed for EMVs without loaded P49, implying that P49 constitutes those particles on the surface of EMVs. These results indicate that P49 is loaded onto EMVs via its interaction with the CPS of EMVs.

**IMPORTANCE:** Elucidating the mechanisms of cargo loading onto bacterial extracellular membrane vesicles (EMVs) is important to understand their biogenesis and to develop their applications. Here, we show that the major cargo protein of EMVs from a fish intestinal Gram-negative bacterium, *Shewanella vesiculosa* HM13, is loaded onto EMVs through its interaction with capsular polysaccharide (CPS) of EMVs. Genes involved in CPS synthesis were also identified. To our knowledge, there have been no reports describing the cargo protein-loading mechanism in which CPS serves as the protein-binding scaffold for EMVs. Thus, this study represents a new mode of protein loading onto EMVs. The results deepen our understanding of cargo loading onto EMVs and would contribute to development of their applications.

## INTRODUCTION

Nano-sized extracellular capsules released from bacteria, called extracellular membrane vesicles (EMVs), have spherical bilayer structures ranging in size from 20 to 250 nm. EMVs play diverse roles in bacterial survival, including intra- and inter-species communication, horizontal gene transfer, pathogenesis, biofilm formation, and self-defense (1, 2). These functions of EMVs depend on their cargo molecules. For instance, EMVs of pathogenic bacteria such as *Pseudomonas aeruginosa*, *Vibrio cholerae*, and enterotoxigenic *Escherichia coli* selectively deliver multiple virulence factors for survival and proliferation within the host, which may result in severe disease (3–5). Protein loading onto EMVs provides cargo proteins with various advantages, including protection from proteolytic degradation, long-distance delivery, host-cell specific targeting, and modulation of the immune response (6). In addition to their physiological importance, vesiculation-mediated protein secretion has potential for various applications, such as the development of drug delivery systems, nanobiocatalysts, vaccines, and heterologous protein production systems (2). Thus, cargo loading onto EMVs is important for their physiological functions and their potential applications.

*Shewanella vesiculosa* HM13, a fish intestinal Gram-negative bacterium, produces abundant EMVs carrying a single major cargo protein named P49 (7). Although the physiological function of P49 is yet to be elucidated and is unpredictable based on its sequence similarity to known proteins, the high purity and quantity of P49 in EMVs raises the expectation that it can be used as a carrier protein to produce a foreign protein as a cargo of EMVs using the recombinant protein production system (7, 8). We previously found that a gene cluster in the neighboring region of the P49 gene is involved in the P49 transport to EMVs (9). This gene cluster contained genes encoding non-canonical type II secretion system (T2SS), homologs of surface polysaccharide-chain synthesis proteins, including a homolog of a flippase involved in bacterial extracellular polysaccharide synthesis, Wzx, and other functionally unknown proteins (**Fig. 1 and Table S1**). Gene disruption analysis indicated that P49 accumulates in the cells in the absence of the non-canonical T2SS, suggesting that this machinery transports P49 to the cell surface or extracellular space. We also found that P49 is secreted to the extracellular space without being loaded onto EMVs in the absence of *hm3343,* coding for Wzx, and other genes in the vicinity of the P49 gene, *hm3342*, *hm3344*, and *hm3345*, suggesting that these genes are essential for the synthesis of surface components of EMVs required for P49 loading onto EMVs (9). However, the validity of this speculation and the identity of the molecule required for P49 loading remain unclear.

**Fig. 1.**
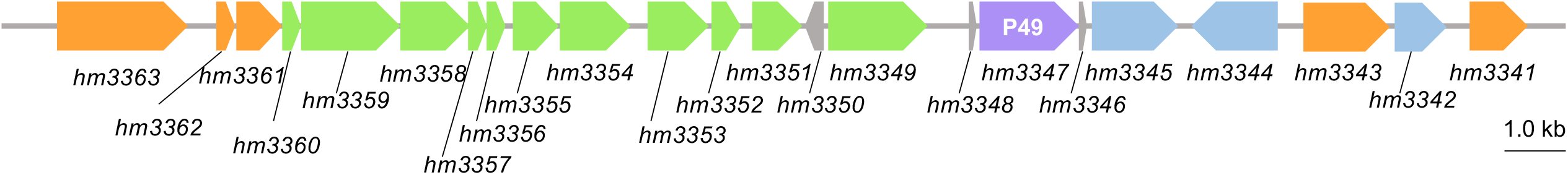
Genetic organization of the P49-gene containing gene cluster. Genetic map of the gene cluster containing the P49 gene and its neighboring genes. Predicted functions and localization of the gene products are listed in **Table S1**. The P49-coding gene is indicated by the purple arrow, while genes having homology to genes involved in polysaccharide synthesis and protein translocation are indicated by orange and green arrows, respectively. Genes coding for homologs of glycerophosphodiester phosphodiesterase, phosphoethanolamine transferase, and nitroreductase are shown by blue arrows.

The cell surface of Gram-negative bacteria is often surrounded by polysaccharide chains, such as the O-antigen of lipopolysaccharide (LPS) and capsular polysaccharide (CPS). Many of them have high antigenic activity and can be involved in cell defense and interaction with their environment (10). Since the surface of EMVs is assumed to mimic the structure of the outer membrane of the cells, surface polysaccharide of EMVs also plays important roles in these processes. Thus, in order to understand the physiological function of the EMVs of *S. vesiculosa* HM13 and to develop their application, it is important to clarify their surface polysaccharide structure. Previous studies determined the chemical structures of LPS and CPS of *S. vesiculosa* HM13 and showed that LPS of this bacterium lacks an O-antigen (11, 12) and that EMVs and cells are covered with the same CPS with a novel aminosugar, named Shewanosamine (13).

In the present study, we investigated the function of HM3343, a Wzx homolog, and other proteins encoded by the genes around the P49 gene in the biosynthesis of the surface polysaccharide chain of *S. vesiculosa* HM13 and in P49 loading onto EMVs. We found that the association of P49 with CPS is the mechanism of P49 loading onto EMVs. This finding provides new insight into protein cargo loading onto bacterial EMVs and contributes to the development of a system for presenting heterologous proteins on EMVs using P49 as a carrier.

## RESULTS

### Wzx-dependent synthesis of EMV-associated CPS

In the P49-gene cluster, the gene designated *hm3343* encodes a putative inner membrane flippase, Wzx (9). Wzx family proteins are generally responsible for the biosynthesis of bacterial cell surface polysaccharides, i.e. CPS and O-antigen of LPS, by mediating the translocation of their oligosaccharide precursors from the cytoplasmic face to the periplasmic face of the inner membrane (14). The chemical structure of LPS from both cells and EMVs of *S. vesiculosa* HM13 was previously determined, indicating that it is rough LPS (LOS), which lacks the O-antigen (11, 12). On the other hand, this strain was found to produce CPS as the surface component of both cells and EMVs (13). Thus, we speculate that the Wzx homolog encoded by *hm3343* is involved in the synthesis of CPS, but not the O-antigen of LPS. In the present study, to verify this hypothesis, EMVs of the parent strain and the *hm3343*-deleted mutant were subjected to CPS analysis following the Tipton and Rather method (15). For the extract of the parent strain (Δ*p49*Δ*pyrF*^HM13^ (**Table S2**)), a smear band of CPS, whose average molecular mass was determined to be 40 kDa (13), was detected in the Alcian blue-stained SDS-PAGE gel in the slow migration zone, whereas this major band was barely visible for the *hm3343*- deleted mutant (Δ*hm3343*Δ*p49*Δ*pyrF*^HM13^ (**Table S2**)) (**Fig. 2A**). The CPS band was detected for the extract from EMVs of Δ*hm3343*Δ*p49*Δ*pyrF*^HM13^ harboring the *hm3343* expression plasmid, p*hm3343* (Δ*hm3343*Δ*p49*Δ*pyrF*^HM13^/p*hm3343* (**Table S2**)) (**Fig. 2A**). The CPS band was also detected for cells of Δ*p49*Δ*pyrF*^HM13^ and Δ*hm3343*Δ*p49*Δ*pyrF*^HM13^ harboring p*hm3343* but not for Δ*hm3343*Δ*p49*Δ*pyrF*^HM13^ cells (**Fig. 2B**). These results indicate that the Wzx homolog, encoded by *hm3343* in the vicinity of the P49 gene, is involved in the synthesis of CPS of both EMVs and cells.

**Fig. 2.**
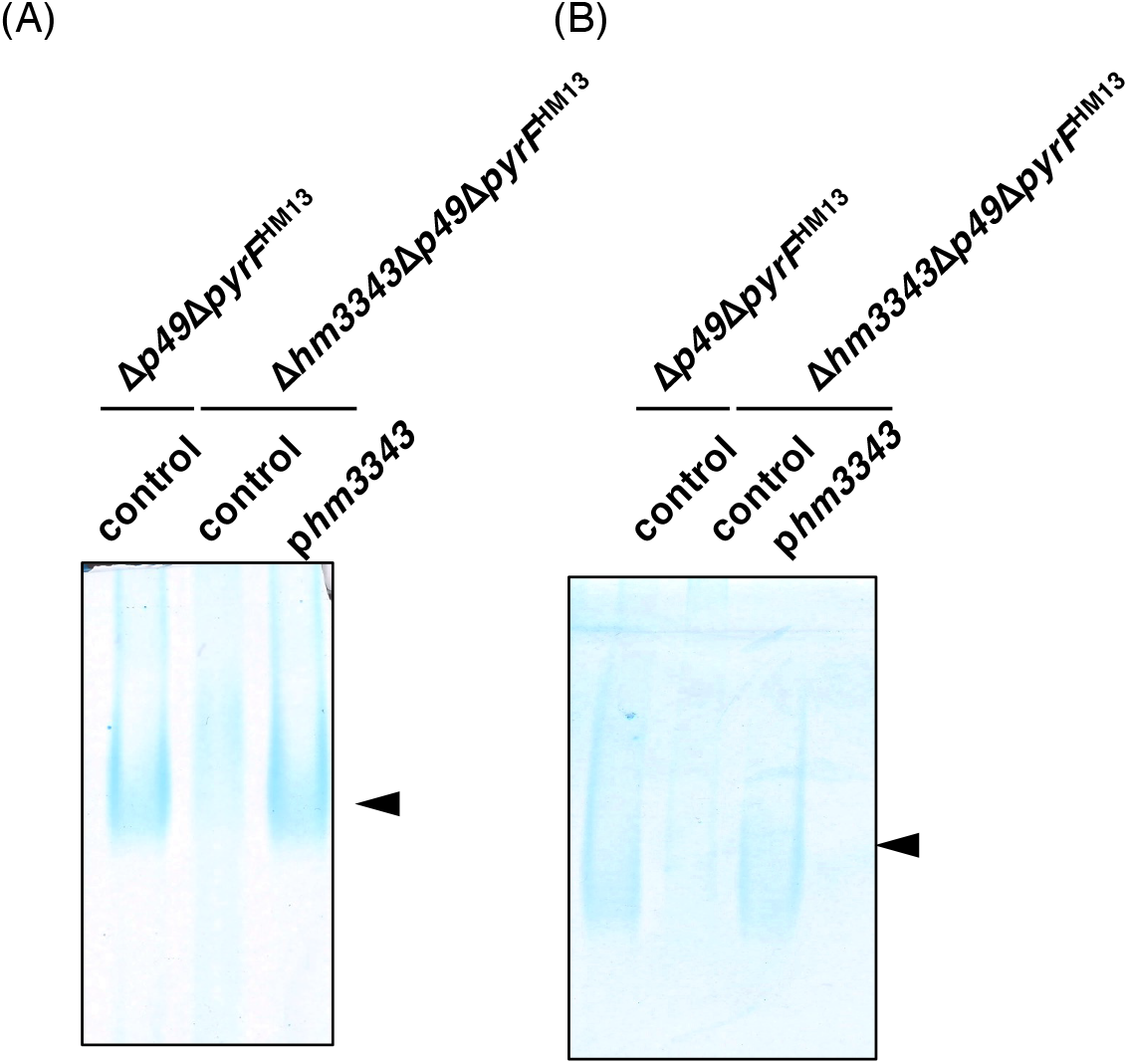
CPS extracted from EMVs and cells of *S. vesiculosa* HM13. Alcian blue-stained SDS-PAGE analysis of CPS extracted from EMVs (A) and cells (B) of Δ*p49*Δ*pyrF*^HM13^ harboring an empty plasmid, pJRD-Cm^r^, (control) and its *hm3343*-deletion mutant, Δ*hm3343*Δ*p49*Δ*pyrF*^HM13^, harboring pJRD-Cm^r^ (control) or the *hm3343*-expression plasmid (p*hm3343*). The arrowheads indicate the positions of CPS.

### CPS-dependent P49 binding to EMVs

To investigate the involvement of CPS in P49 loading onto EMVs, we conducted an *in vitro* P49-binding assay using purified P49 and P49-free EMVs prepared from the P49-less mutant (Δ*p49* (**Table S2**)). P49 was purified from the post-vesicle fraction (PVF), an EMV-free culture supernatant obtained by pelleting EMVs, from a mutant that lacks *hm3345* encoding glycerophosphodiester phosphodiesterase homolog (DDBJ: LC431025) (Δ*hm3345* (**Table S2**)), in which P49 accumulates in PVF without being associated with EMVs (9). The *in vitro* binding assay was conducted using different concentrations of purified P49 (3, 1.5, and 0.75 μM). Purified P49 (100 μL) was mixed with an equal volume of EMVs (equivalent to the amount present in 5 mL culture) to final concentrations of 1.5, 0.75, and 0.375 μM, respectively, and incubated for 15 h. As shown in **Fig. 3A**, purified P49 was observed in the pellet fraction in the presence of EMVs, and the amount of precipitated P49 increased in accordance with increased input P49. In contrast, in the absence of EMVs, the P49 band was detected only in the supernatant. These results indicate that P49 bound to EMVs *in vitro* and co-precipitated with EMVs. It should be noted that the band intensities detected for the EMV pellet were weaker than those for the supernatant of the EMV-free sample. This may be due to degradation of P49 by proteases possibly present in the EMV sample.

**Fig. 3.**
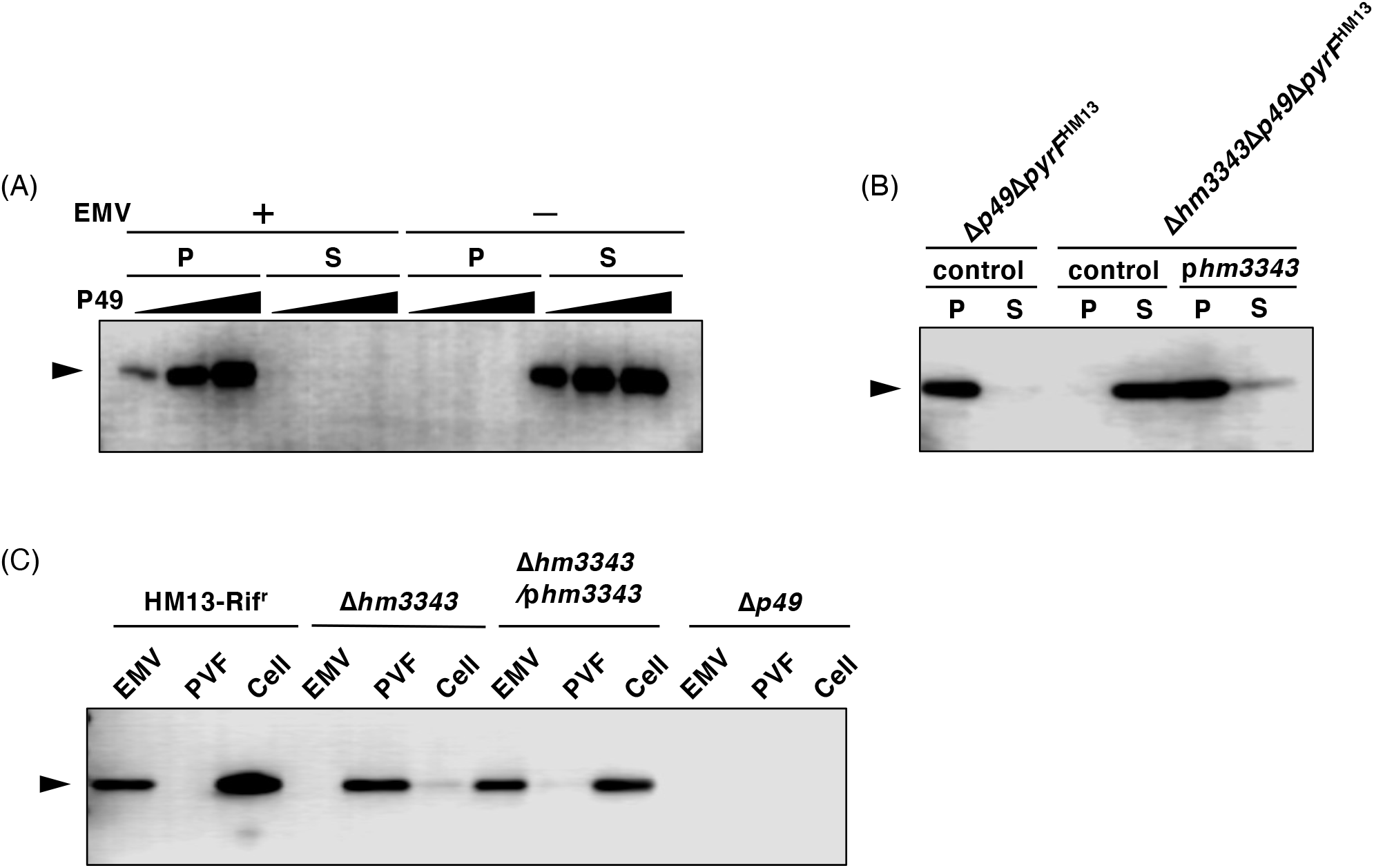
I*n vitro* and *in vivo* P49 binding to EMVs. (A) *In vitro* binding of P49 to P49-free EMVs of Δ*p49*Δ*pyrF*^HM13^. After incubation, samples were fractionated by ultracentrifugation, and the supernatant and pellet fractions were subjected to western blotting with an antibody against P49. As a control, purified P49 without EMVs was subjected to the analysis. (B) *In vitro* P49 binding to EMVs from Δ*p49*Δ*pyrF*^HM13^ harboring an empty plasmid, pJRD-Cm^r^, (control) and *hm3343*-deleted mutant, Δ*hm3343*Δ*p49*Δ*pyrF*^HM13^, harboring pJRD-Cm^r^ (control) or the expression plasmid of *hm3343* (p*hm3343*). P and S indicate pellet and supernatant fractions, respectively. (C) *In vivo* localization of P49 was analyzed for the parent (HM13-Rif^r^), Δ*hm3343*, and Δ*hm3343* harboring the *hm3343*-expression plasmid (Δ*hm3343*/p*hm3343*). P49 in the EMVs, PVF, and cells was detected by western blotting with an antibody against P49. The amount of each sample corresponded to that obtained from 50 µL of culture. The position of P49 is indicated by the arrowhead. These results were confirmed in three independent experiments.

To investigate the effect of the lack of CPS on the interaction between P49 and EMVs, we conducted the *in vitro* binding assay using P49 and EMVs from the CPS-less mutant (Δ*hm3343*Δ*p49*Δ*pyrF*^HM13^ (**Table S2**)). In this assay, the final concentration of P49 was 2 μM. As a result, P49 was detected in the supernatant (**Fig. 3B**). The ability of EMVs to interact with P49 was restored by introducing a *hm3343*-complementation plasmid, p*hm3343* (Δ*hm3343*Δ*p49*Δ*pyrF*^HM13^/p*hm3343* (**Table S2**)). The dissociation of P49 from EMVs was also observed *in vivo* for Δ*hm3343* (**Table S2**) (**Fig. 3C**). The P49-loading capacity of the EMVs of this strain was restored by the introduction of p*hm3343* (Δ*hm3343*/p*hm3343* (**Table S2**)) (**Fig. 3C**). These results demonstrated that CPS synthesized by the HM3343-dependent pathway plays a crucial role in the association between P49 and EMVs both *in vivo* and *in vitro*.

### Occurrence of P49 as peripheral particles around EMVs

We analyzed the morphology of EMVs from the parent strain (Δ*pyrF*^HM13^) and Δ*p49*Δ*pyrF*^HM13^ and EMVs from Δ*p49*Δ*pyrF*^HM13^ incubated with P49. Transmission electron microscopy (TEM) analysis of the parent EMVs demonstrated the presence of peripheral particles with a diameter of 10.2 +/- 3.0 nm (n = 100) on the surface of spherical vesicles with a diameter of 35.0 +/- 3.2 nm (n = 100) (**Fig. 4A**). Peripheral particles were not observed around the EMVs of the P49-less mutant (**Fig. 4B**). Peripheral particles with a size of 9.5 +/- 3.0 nm (n = 100), similar to those observed for the EMVs of the parent strain, were observed around the EMVs after *in vitro* binding of purified P49 (**Fig. 4C**). These results suggest that P49 constitutes peripheral particles of the EMVs.

**Fig. 4.**
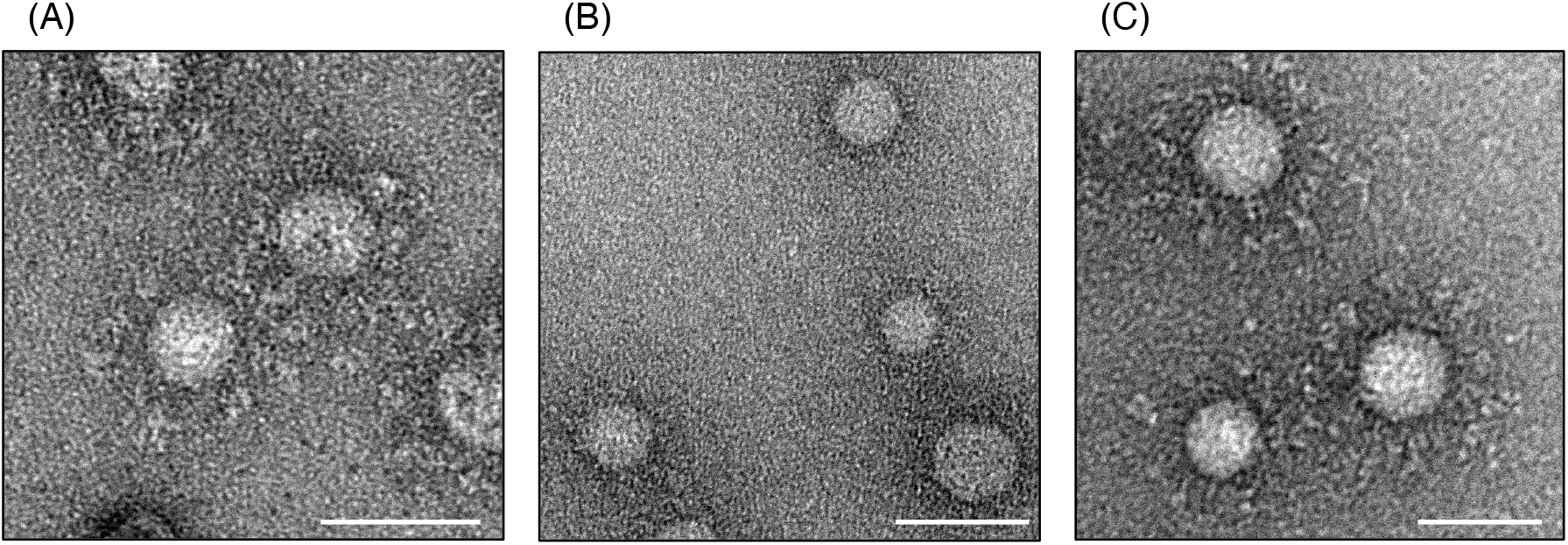
TEM analysis of EMVs with and without loaded P49. TEM images of EMVs from Δ*pyrF*^HM13^ (A) and Δ*p49*Δ*pyrF*^HM13^ (B) and EMVs from Δ*p49*Δ*pyrF*^HM13^ incubated with purified P49 *in vitro* (C). Scale bars indicate 50 nm.

### Characterization of the *in vitro* association between P49 and EMVs

To further characterize the *in vitro* association between P49 and EMVs, we analyzed the secondary structure of EMV-free P49 and P49 associated with EMVs of Δ*p49* (**Table S2**) using circular dichroism (CD) analysis (**Fig. 5A**). We observed that the purified P49 did not exhibit a typical CD signal of proteins with α-helix and β-sheet structures. In contrast, P49 incubated with EMVs from Δ*p49* showed a typical CD signal of β-sheet proteins, with the lowest negative ellipticity around 215 nm. The structural change in P49 caused by incubation with the EMVs from Δ*p49* was not detected when the EMVs of Δ*hm3343* (**Table S2**), which did not bind to P49 as shown in **Fig. 3C**, were used.

**Fig. 5.**
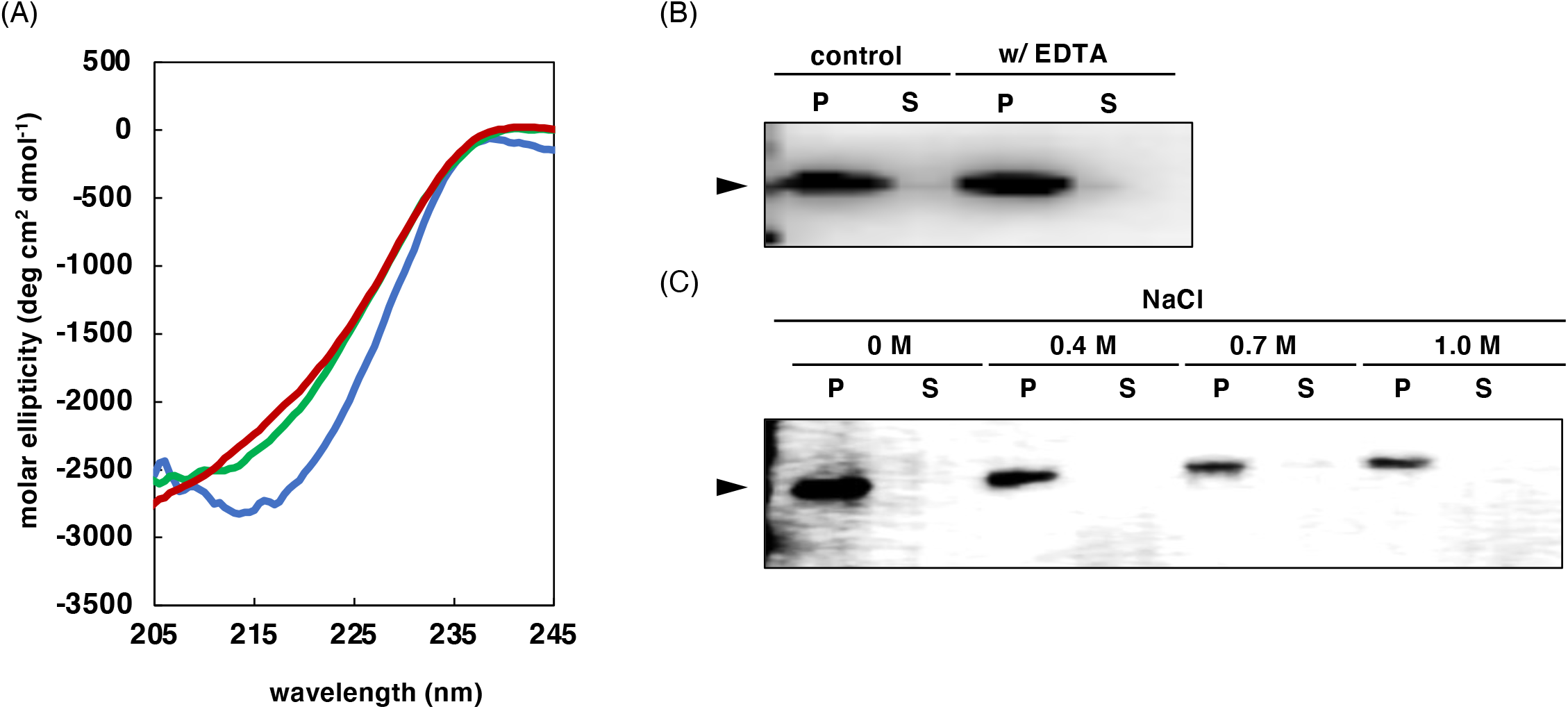
Characterization of the *in vitro* P49 binding to EMVs. (A) The CD spectra of the EMV-free purified P49 (red) and P49 incubated with the EMVs of Δ*p49* (blue) and Δ*hm3343* (green). (B) Effects of EDTA supplementation on *in vitro* P49 binding to EMVs. *In vitro* P49 binding to EMVs was performed in the presence (w/ EDTA) and absence (control) of 20 mM EDTA. (C) *In vitro* P49 binding to EMVs in the presence of various concentrations of NaCl (0, 0.4, 0.7, and 1.0 M). P and S indicate pellet and supernatant fractions, respectively. Three independent experiments confirmed the results.

To examine whether divalent metal ions such as Ca^2^ ^+^ and Mg^2+^ play a role in the interaction between P49 and EMVs, the cargo loading assay using P49 and EMVs of Δ*p49*Δ*pyrF*^HM13^ was conducted in the presence of a chelating agent, ethylenediaminetetraacetic acid (EDTA). In the binding assay, the final concentration of P49 was 2 μM. As shown in **Fig. 5B**, supplementation with 20 mM EDTA did not affect the association of P49 with EMVs, and no protein band was observed in the supernatant. Thus, it is unlikely that divalent metal ions are involved in the binding of P49 to EMVs.

We also analyzed the effects of ionic strength on the interaction between P49 and EMVs by increasing the NaCl concentration in the *in vitro* P49-loading assay mixture. We observed that P49 was co-precipitated with EMVs even in the presence of 1.0 M NaCl (**Fig. 5C**), suggesting that electrostatic interactions do not play an important role in the interaction between P49 and EMVs.

### Involvement of P49-neigboring genes in CPS synthesis and P49 loading onto EMVs

In the vicinity of the P49 gene, we identified genes encoding the subunits of a non-canonical T2SS [*hm3360* (*gspG2,* major pseudopilin), *hm3359* (*gspE2,* secretion ATPase), *hm3358* (*gspF2*, inner membrane platform), *hm3354* (*gspK2,* minor pseudopilin), *hm3351* (*gspB2*, accessory protein), and *hm3349* (*gspD2*, outer membrane conduit)] and genes encoding homologs of polysaccharide export protein (*hm3363*), undecaprenyl-phosphate alpha-*N*-acetylglucosaminyl 1-phosphate transferase (*hm3362* and *hm3361*), glycerophosphodiester phosphodiesterase (*hm3345*), phosphoethanolamine transferase (*hm3344*), Wzx flippase (*hm3343*), nitroreductase (*hm3342*), and UDP-glucose 6-dehydrogenase (*hm3341*) (7) (**Fig. 1 and Table S1**). To examine whether these gene products are involved in CPS synthesis, the EMVs of the mutants obtained by disrupting *hm3359*, *hm3358*, *hm3354*, *hm3349*, *hm3345*, *hm3344*, *hm3343*, and *hm3342* individually were subjected to CPS analysis. As the result, the broad band observed in the low-mobility area for the parent strain was barely visible for

Δ*hm3345*, Δ*hm3344*, and Δ*hm3343* (**Fig. 6A**). In contrast, the CPS band was detected in EMVs from the mutants that lacked the non-canonical T2SS components (Δ*hm3359*, Δ*hm3358*, Δ*hm3354*, and Δ*hm3349*) and the nitroreductase homolog (Δ*hm3342*). To examine whether P49 was loaded onto EMVs from these mutants, an *in vitro* P49-binding assay was conducted. In this assay, P49 was added at a final concentration of 2 μM, and the mixture was incubated for 1 h at 4 °C. EMVs from the mutants that lacked the non-canonical T2SS components showed binding capacity for purified P49, whereas the disruption of *hm3345*, *hm3344*, *hm3343*, and *hm3342* impaired P49 loading onto EMVs (**Fig. 6B**).

**Fig. 6.**
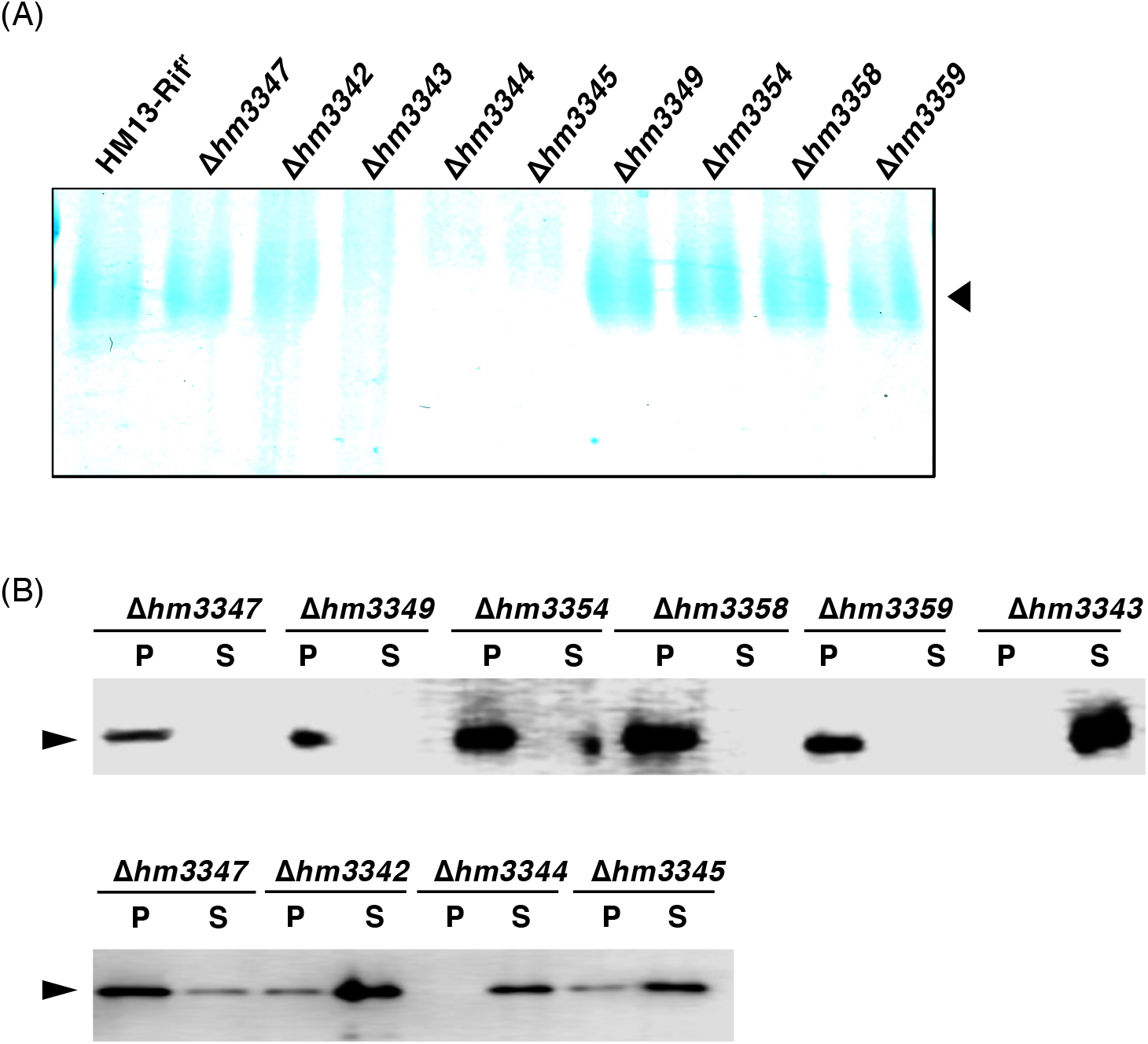
Effects of the disruption of the genes around the P49 gene on the CPS and the P49-binding capacity of the EMVs. (A) Analysis of the CPS extracts of EMVs prepared from the parent strain and gene-disrupted mutants. The arrowhead indicates the position of CPS. (B) *In vitro* P49-binding assay using EMVs from the parent and mutants. After *in vitro* P49 binding, the samples were separated into pellet (P) and supernatant (S) fractions by ultracentrifugation. The arrowheads indicate the position of P49.

## DISCUSSION

Here, we identified the genes involved in the synthesis of EMV-associated CPS of *S. vesiculosa* HM13, whose chemical structure was recently determined and shown to be identical to that of cellular CPS (13) (**Figs. 2 and 6A**). By disrupting these genes, we revealed that P49, the major cargo protein of EMVs, is bound to EMVs in a CPS-dependent manner. To demonstrate the CPS-dependent binding of P49 to EMVs, we conducted an *in vitro* P49-binding assay. P49 bound to EMVs from the parent strain producing CPS, whereas P49 did not bind to EMVs from mutants that did not produce CPS (**Figs. 3B and 6B**). We observed small particles around the EMVs from the parent strain and those from the P49-less mutant incubated with P49 *in vitro* by TEM analysis, whereas such particles were not observed around EMVs not carrying P49 (**Fig. 4**). These results imply that P49 is loaded onto EMVs via its interaction with CPS on the surface of the EMVs (**Fig. 7**). CD analysis suggested that P49 adopted a β-strand-rich structure upon its binding to EMVs (**Fig. 5A**). Considering the size of the peripheral particles of EMVs, it is reasonable to assume that the particles consist of P49 oligomers or P49 complexed with CPS (**Fig. 4**). Previous studies showed that P49 accumulates in the cells of gene-disrupted mutants that lack the T2SS-like machinery encoded by the genes upstream of the P49 gene (*hm3359*, *hm3358*, *hm3354*, and *hm3349*) (9), indicating that this machinery is responsible for the translocation of P49 across the outer membrane (**Fig. 7**).

**Fig. 7.**
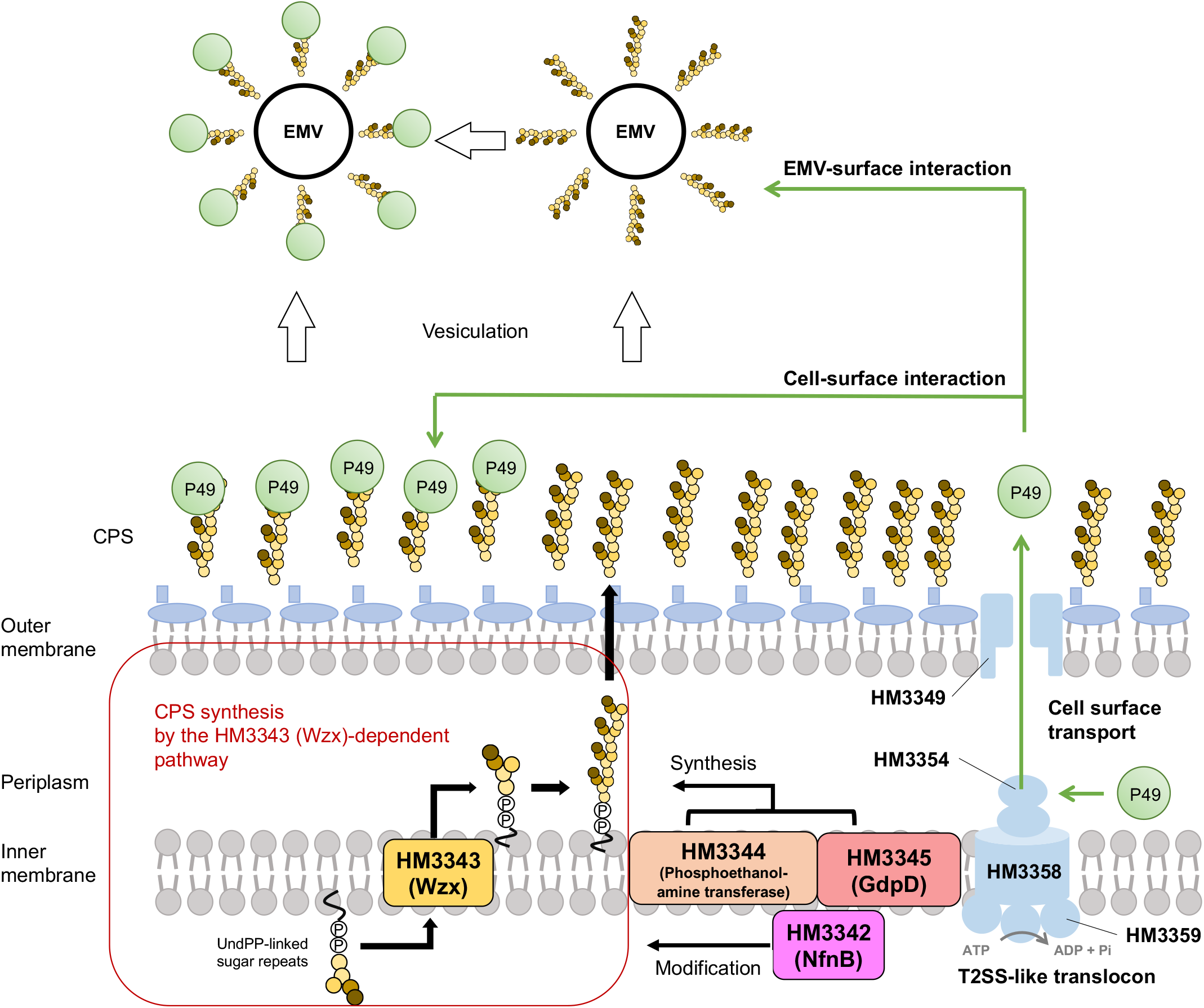
The model of P49 transport onto EMVs of *S. vesiculosa* HM13. P49 is translocated across the outer membrane through the T2SS-like translocon and is subsequently loaded onto EMVs through its interaction with CPS. In CPS synthesis, Wzx encoded by *hm3343* mediates the translocation of undecaprenyl pyrophosphate (UndPP)-linked sugar repeats across the inner membrane from the cytoplasm to the periplasmic space. Homologs of phosphoethanolamine transferase (HM3344) and GdpD (HM3345) are also required for CPS synthesis. NfnB (HM3342) is probably responsible for the CPS modification required for the interaction with P49. Protein localization was predicted using the PSORTb version 3.0.3 (http://www.psort.org/psortb/).

We previously found that P49 is released into the culture supernatant without being loaded onto EMVs upon disruption of genes coding for homologs of Wzx (HM3343), phosphoethanolamine transferase (HM3344), and glycerophosphodiester phosphodiesterase (HM3345), located downstream of the P49 gene (9). However, it was unclear why P49 was not loaded onto the EMVs in these mutants. In the present study, we demonstrated that this was due to the lack of CPS on the surface of the EMVs in these mutants. We found that HM3343, HM3344, and HM3345 play a pivotal role in CPS synthesis (**Figs. 2, 6A, and 7**). The primary structure of HM3343 strongly suggests that it functions as a flippase to translocate CPS precursors across the inner membrane (14, 16). However, it is not clear how HM3344 and HM3345 are involved in CPS synthesis based on *in silico* analysis. Although HM3344 has sequence similarity to phosphoethanolamine transferase (17, 18), no phosphoethanolamine residue was found in CPS (13). As to HM3345, most homologs of this protein are registered as hypothetical proteins in databases. Although some homologs are annotated as glycerophosphodiester phosphodiesterase (19, 20), their relevance to CPS synthesis has not been reported. Thus, their function in CPS synthesis should be examined experimentally in future studies. Notably, the disruption of *hm3342* adjacent to *hm3343* (*wzx*) did not cause the loss of the CPS band in SDS-PAGE analysis (**Fig. 6A**) but resulted in the loss of interaction between EMVs and P49 (**Fig. 6B**). Accordingly, we consider that *hm3342* is not required for the polymerization step in CPS biosynthesis but is required for the generation of a CPS moiety that is essential for the interaction between CPS and P49 (**Fig. 7**). Since HM3342 shows sequence similarity with nitroreductase (21–23), which is a redox enzyme, it is plausible to assume that it catalyzes a redox reaction in CPS biosynthesis to generate a structural moiety of CPS for its interaction with P49. A comparison between CPS from Δ*hm3342* and the parent strain will help to uncover this structural moiety of CPS. In addition to *hm3343*, *hm3344*, *hm3345*, and *hm3342*, many genes in the vicinity of the P49 gene are speculated to be involved in CPS biosynthesis (**Fig. 1** and **Table S1**), such as genes encoding glycosyltransferase homologs (*hm3367, hm3340, hm3338*, and *hm3311*). Because the CPS of this strain has a novel structure with a unique amino sugar, which has never been reported before (13), CPS biosynthetic enzymes with unique catalytic properties are expected to be discovered.

It is noteworthy that CPS serves as a tethering scaffold for protein cargo loading onto EMVs. Previous studies revealed that proteins with a C-terminal signal for the Type IX secretion system are conjugated to anionic LPS as a membrane anchor and enriched in EMVs of *Porphyromonas gingivalis* (24, 25). Other studies showed that surface-exposed lipoproteins with a lipoprotein export sequence and with acyl groups introduced at the N-terminus are enriched in EMVs of *Bacteroides* (26, 27). These studies revealed that anionic LPS and N-terminal acyl groups tether proteins to EMVs. However, there have been no reports, to our knowledge, describing the occurrence of cargo proteins loaded onto EMVs through their interaction with CPS. Thus, this study represents a new mode of protein loading onto EMVs.

The results obtained in this study are also significant as they pave the way for the use of CPS as a scaffold for protein display on the surface layer of EMVs. Recombinant protein display on EMVs is expected to be a useful technique for the development of nanoscale materials. For example, vaccines and therapeutic agents have been developed by displaying recombinant proteins on bacterial EMVs (28–30). Currently available methods for loading foreign proteins onto the surface of EMVs employ membrane anchor proteins as the fusion/binding partners of foreign proteins (30, 31). The interaction between P49 and CPS discovered in this study may offer a new method to display foreign proteins on the surface of EMVs: foreign proteins fused with P49 may be displayed on the surface of EMVs through their interaction with CPS present on the outside of the membrane of EMVs. This method is different from conventional methods in that foreign proteins can be displayed through their interaction with CPS without using membrane components, such as membrane proteins and LPS, as anchors to EMVs. Modifying membrane components may have undesirable effects on membrane dynamics and stability. The P49-CPS system may be useful in avoiding such adverse effects. Additionally, this system would allow protein loading onto EMVs after vesiculation completion, as expected by the fact that P49 was loaded onto isolated EMVs *in vitro*. This may expand the variation of foreign proteins loaded onto EMVs: proteins that are difficult to produce by *S. vesiculosa* HM13 may be produced by other cells and then loaded onto the EMVs of *S. vesiculosa* HM13.

Although it is a future subject to establish a system for foreign protein loading onto EMVs using CPS as a binding scaffold, this study is significant in that it revealed a new protein-loading mechanism onto EMVs and demonstrated the possibility of using CPS as a scaffold for protein loading onto EMVs. Detailed mechanistic elucidation of the interaction between P49 and CPS is also a subject for future research. Determination of the structural moieties of P49 and CPS involved in their interaction is important for developing the protein-loading systems described above. The *in vitro* cargo-loading system established in this study facilitates such mechanistic analyses.

## MATERIALS AND METHODS

### Bacterial strains and growth conditions

The strains used in this study are listed in **Table S2**. The strains of *S. vesiculosa* HM13 were cultured in 30 mL LB liquid medium at 18 °C in a Bio Shaker BR-43FL incubator (Taitec, Saitama, Japan) at 180 rpm up to OD_600_ = 2–3. *Escherichia coli* S17- 1/λ*pir* (32) used as a plasmid donor cell for the *pir*-dependent gene knockout plasmid, pKKPH, was grown in 5 mL LB medium at 37 °C. When required, uracil, 5-fluoroorotic acid (5-FOA), kanamycin (Km), rifampicin (Rif), and chloramphenicol (Cm) were added to the medium at final concentrations of 40 µg/mL, 1 mg/mL, 50 µg/mL, 50 µg/mL, and 30 µg/mL, respectively.

### Gene deletion and complementation in *S. vesiculosa* HM13

We obtained the gene-deleted and -inserted mutants based on a previously described method using the gene coding for orotidine-5′-phosphate decarboxylase (*pyrF*) (33). Based on the whole-genome sequence, we identified a *pyrF*-coding gene for *S. vesiculosa* HM13 (*pyrF*^HM13^). The construction method of the *pyrF*^HM13^-deleted strain (Δ*pyrF*^HM13^) (**Table S2**) and pKKPH plasmid for *pyrF*-based gene knockout is described in Supplemental procedures. Δ*p49*Δ*pyrF*^HM13^ and Δ*hm3343*Δ*p49*Δ*pyrF*^HM13^ were constructed using the following two-step selection. The primers used are listed in **Table S3**. The upstream and downstream regions of the P49 gene and *hm3343* were amplified from the genomic DNA of *S. vesiculosa* HM13 by PCR with the following primer pairs: *p49*_upfwd/*p49*_uprev, *p49_*downfwd/*p49_*downrev, *wzx*_upfwd/*wzx*_uprev, and *wzx*_downfwd/*wzx*_downrev, respectively. A linear fragment of the knockout plasmid pKKPH was amplified using pKKPH-invfwd and pKKPH-invrev. To generate the *pyrF*^HM13^-based gene-deletion plasmids (pKKPH-*p49* and pKKPH-*hm3343*), each of the upstream and downstream fragments was ligated with the pKKPH fragment using the NEBuilder HiFi DNA Assembly Cloning Kit (New England Biolabs, Japan, Tokyo, Japan). To insert pKKPH-*p49* to the genome of Δ*pyrF*^HM13^, pKKPH-*p49* was introduced into *E. coli* S17-1/*λpir* and transferred to Δ*pyrF*^HM13^ by conjugation. Transformants were selected on LB plates containing uracil, Rif, and Km as the first selection. The selected mutant cells were then transferred to LB plates containing uracil, Rif, and 5-FOA to isolate 5-FOA-resistant single colonies as the second selection. The *p49*-deleted mutants were selected by diagnostic PCR using the following set of primers: Check_*p49*fwd/Check_*p49*rev. To obtain a double mutant lacking both *p49* and *wzx,* pKKPH-*hm3343* was introduced into Δ*p49*Δ*pyrF*^HM13^, and the *hm3343*-deleted mutant (Δ*hm3343*Δ*p49*Δ*pyrF*^HM13^) was selected via two-step selection, as described above. The deletion of *hm3343* was confirmed by PCR using Check_*wzx*fwd/Check_*wzx*rev.

For *hm3343*-complementation, an expression plasmid carrying *hm3343* (p*hm3343*) was constructed. Linearized pJRD-Cm^r^ was amplified by PCR using the following primer set: pJRD-invfwd/pJRD-invrev. The DNA fragment of *hm3343* and the 474 bp upstream flanking region of *hm3343* containing its predicted promoter was amplified with the primer set *wzx*up500fwd/*wzx*rev. PCR products were assembled using the NEBuilder HiFi DNA Assembly Cloning Kit to obtain p*hm3343*. The *wzx* insertion was confirmed by DNA sequencing of the plasmid using the primers pJRD_CheckFW1, *wzx*_CheckFw2, and pJRD_CheckRV. The constructed plasmid was introduced into *E. coli* S17-1/*λpir* cells and transferred to Δ*hm3343*Δ*p49*Δ*pyrF*^HM13^ and Δ*hm3343* by conjugation. Δ*hm3343*Δ*p49*Δ*pyrF*^HM13^ and *Δhm3343* harboring p*hm3343* (Δ*hm3343*Δ*p49*Δ*pyrF*^HM13^/p*hm3343* and Δ*hm3343/*p*hm3343*, respectively) were selected on LB plates containing Rif and Cm. As controls, empty pJRD-Cm^r^ was introduced into Δ*p49*Δ*pyrF* ^HM13^ and Δ*hm3343*Δ*p49*Δ*pyrF*^HM13^ to generate Δ*p49*Δ*pyrF* ^HM13^*/*p and Δ*hm3343*Δ*p49*Δ*pyrF*^HM13^/p, respectively.

### Isolation of EMVs

EMVs were isolated using a previously reported method (9, 34). Briefly, cells were pelleted by centrifugation at 6,000 ×*g* at 4 °C for 15 min with ALLEGRA X-30R (Beckman Coulter, Brea, CA, USA). The supernatant was centrifuged at 15,000 ×*g* at 4 °C for 15 min with Sorvall LYNX4000 (Thermo Scientific, Waltham, MA, USA). To remove the remaining bacterial cells, the supernatant was filtered through a 0.45-µm pore filter. The filtrate was ultracentrifuged at 100,000 ×*g* at 4 °C for 2 h using an Optima XE-90 (Beckman Coulter) to collect EMVs. The EMV pellets were suspended in 500 µL of Dulbecco’s phosphate buffered saline (135.9 mM NaCl, 2.7 mM KCl, 8.9 mM Na_2_HPO_4_, and 1.5 mM KH_2_PO_4_, pH7.2) supplemented with 0.2 M NaCl (DPBSS). To quantify the EMVs, the collected EMVs were stained with a lipophilic fluorescent molecule, *N*-(3-triethylammoniumpropyl)-4-(*p*-diethylaminophenyl-hexatrienyl) pyridinium dibromide (FM4-64) (Thermo Fisher Scientific, Waltham, MA, USA) as previously described (9). Based on the results, the same amount of EMVs was used for the CPS analysis and *in vitro* P49 binding assay for different samples.

### Purification of P49

P49 for *in vitro* binding with EMVs was prepared from 200 mL culture supernatant of Δ*hm3345*, which secretes P49 into the extracellular space without association with EMVs. After removing the bacterial cells, the supernatant was concentrated with an Amicon Ultra filter (Merck Millipore, Billerica, MA, USA; 30 kDa molecular weight cut-off) and subjected to ultracentrifugation at 100,000 ×*g* for 2 h at 4 °C to remove EMVs. The supernatant (post-vesicle fraction, PVF) was then fractionated with Superose 12 10/300 GL column (GE Healthcare, Little Chalfont, UK) (flow rate 0.4 mL/min, fraction volume 2.0 mL). Fractions were eluted with PBS buffer (137 mM NaCl, 2.7 mM KCl, 10 mM Na_2_HPO_4_·12H_2_O, and 1.8 mM KH_2_PO_4_, pH 7.4). The purity of P49 in the eluted fractions was analyzed by SDS-PAGE and staining with Coomassie Brilliant Blue G-250. The concentration of P49 was measured using a BCA protein assay kit (Nacalai Tesque, Kyoto, Japan) with bovine serum albumin as a standard.

### Extraction and characterization of CPS

CPS was extracted from the EMVs using a previously reported method (15) with some modifications. Briefly, 200 µL of CPS extraction buffer (60 mM Tris-HCl (pH 8.0), 0.045 mM CaCl_2_, and 21 mM MgCl_2_) was added to 100 µL of the EMV suspension, which contained EMVs corresponding to 5 mL culture. The mixture was subjected to three freeze-thaw cycles at -80 °C and 37 °C and treated with 5 units of DNase I (TaKaRa Bio, Otsu, Japan) and 20 ng RNase (Nippon Gene, Tokyo, Japan) at 37 °C for 30 min to remove nucleic acids. The reaction was stopped by the addition of SDS to a final concentration of 0.33% (w/v) and boiling. The mixture was then treated with 300 µg proteinase K (Thermo Scientific) at 60 °C for 1 h to remove proteins and centrifuged to remove debris. Finally, CPS was precipitated overnight in 75% ice-cold ethanol. The CPS pellet was dissolved in SDS-PAGE loading buffer (125 mM Tris-HCl (pH 6.8), 10% (v/v) 2-mercaptoethanol, 4% (w/v) SDS, 10% (w/v) sucrose, and 0.01% (w/v) bromophenol blue) and heated at 100 °C for 5 min. CPS was visualized by 12.5% SDS-PAGE and staining with Alcian Blue Solution (Wako Pure Chemical Industries, Osaka, Japan).

### *In vitro* P49 binding assay

Purified P49 and EMVs were mixed at the ratios and conditions shown in the Results section. The mixtures were incubated at 4 °C. Incubation times are described in the Results section. After incubation, the P49-EMV mixtures were ultracentrifuged at 100,000 ×*g* for 2 h at 4 °C using an Optima XE-90 to collect the EMV pellet and supernatant. The pellet and supernatant were subjected to trichloroacetic acid precipitation and analyzed by 12.5% SDS-PAGE and western blotting with an anti-P49 antibody.

### TEM analysis

The TEM images of the EMVs were obtained as follows. The EMV samples (2.5 µL) applied to hydrophilized carbon-coated copper grids were treated twice with 0.5% uranyl acetate. A JEM-1400 transmission electron microscope (JEOL, Tokyo, Japan) at an accelerating voltage of 120 kV and a charge-coupled device (CCD) camera (built-in camera within JEM-1400) were used to obtain images. The diameters of EMVs and peripheral particles around EMVs were measured using the image analysis software Fiji (ImageJ2 version 2.3.0/1.53f) (35).

### CD spectroscopy

CD spectra of EMV-bound and free P49 were obtained using a cuvette with a light path length of 10 mm and a J-820 CD spectrometer (JASCO, Tokyo, Japan). The far-UV spectrum was recorded in the spectral range 200–250 nm at 18 °C.

## Supporting information

Supplemental procedures

## ACKNOWLEDGMENTS

TEM observations were performed in collaboration with the Analysis and Development System for Advanced Materials (ADAM) at the Research Institute for Sustainable Humanosphere, Kyoto University.

