## Supplementary material for "Capsular polysaccharide-mediated protein loading onto extracellular membrane vesicles of a fish intestinal bacterium, *Shewanella vesiculosa* HM13": Table S1

**Table S1 Predicted functions and localization of proteins encoded by the genes in the P49 gene cluster**

| Gene | Function | Localization** | Accession |
|---|---|---|---|
| *hm3363* | Wza, EpsE, polysaccharide export protein | PE | LC431043 |
| *hm3362* | WecA, undecaprenyl-phosphate alpha-*N*-acetylglucosaminyl 1-phosphate transferase | ND | LC431042 |
| *hm3361* | WecA, undecaprenyl-phosphate alpha-*N*-acetylglucosaminyl 1-phosphate transferase | IM | LC431041 |
| *hm3360* | GspG2, Type II secretion system, major pseudopilin | IM | LC431040 |
| *hm3359* | GspE2, Type II secretion system, secretion ATPase | CY | LC431039 |
| *hm3358* | GspF2, Type II secretion system, inner membrane platform protein | IM | LC431038 |
| *hm3357* | Pilin-like protein | ND | LC431037 |
| *hm3356* | Pilin-like protein | ND | LC431036 |
| *hm3355* | Pilin-like protein | ND | LC431035 |
| *hm3354* | GspK2, Type II secretion system, minor pseudopilin | CY | LC431034 |
| *hm3353* | PilN-like protein*, Type IV pilus assembly protein | ND | LC431033 |
| *hm3352* | PilO-like protein*, Type IV pilus inner membrane component | ND | LC431032 |
| *hm3351* | GspB2*, Type II secretion system, accessory protein | IM | LC431031 |
| *hm3350* | Hypothetical protein, no significant similarity found | ND | LC431030 |
| *hm3349* | GspD2, Type II secretion outer membrane pore forming protein, secretin | OM | LC431029 |
| *hm3348* | Hypothetical protein, no significant similarity found | ND | LC431028 |
| *hm3347* | P49, a major cargo protein of EMVs | EX | LC431027 |
| *hm3346* | Hypothetical protein, no putative conserved domains | ND | LC431026 |
| *hm3345* | GdpD, glycerophosphodiester phosphodiesterase | IM | LC431025 |
| *hm3344* | Phosphoethanolamine transferase | IM | LC431024 |
| *hm3343* | Wzx, a flippase to translocate extracellular polysaccharide precursors across inner membrane | IM | LC431023 |
| *hm3342* | NfnB, nitroreductase family protein | CY | LC431022 |
| *hm3341* | Ugd, UDP-glucose 6-dehydrogenase, provisional | CY | LC431021 |

* Estimated function based on structural prediction programs, I-TASSER (https://zhanggroup.org/I-TASSER/) and AlphaFold2 (https://colab.research.google.com/github/sokrypton/ColabFold/blob/main/AlphaFold2.ipynb)
** Protein localization was predicted with PSORTb version 3.0.3 (https://www.psort.org/psortb/). EX, Extracellular space; OM, outer membrane; PE, Periplasm; IM, inner membrane; CY, cytoplasm; ND, not determined
