## Supplementary material for "Capsular polysaccharide-mediated protein loading onto extracellular membrane vesicles of a fish intestinal bacterium, *Shewanella vesiculosa* HM13": Table S2

### Table S2 Strains and plasmids used in this study

| | Description | Reference |
|---|---|---|
| ***Shewanella vesiculosa*** | | |
| HM13-Rif$^r$ | Rifampicin-resistant mutant of *S. vesiculosa* HM13, the parent strain of Δ*pyrF*$^{HM13}$. | (1) |
| Δ*pyrF*$^{HM13}$ | Gene deletion mutant of *pyrF*$^{HM13}$ (*hm2280*), a parent strain of gene deletion mutants | This study |
| Δ*p49*Δ*pyrF*$^{HM13}$ | Deletion mutant of the gene coding for P49 (*hm3347,* LC431027) of Δ*pyrF*$^{HM13}$ | This study |
| Δ*p49*Δ*pyrF*$^{HM13}$/p | Δ*p49*Δ*pyrF*$^{HM13}$ harboring pJRD-Cm$^r$ | This study |
| Δ*hm3343*Δ*p49*Δ*pyrF*$^{HM13}$ | *hm3343* (*wzx,* LC431023)-deletion mutant of Δ*p49*Δ*pyrF*$^{HM13}$ | This study |
| Δ*hm3343*Δ*p49*Δ*pyrF*$^{HM13}$/p | Δ*hm3343*Δ*p49*Δ*pyrF*$^{HM13}$ harboring pJRD-Cm$^r$ | This study |
| Δ*hm3343*Δ*p49*Δ*pyrF*$^{HM13}$/p*hm3343* | Δ*hm3343*Δ*p49*Δ*pyrF*$^{HM13}$ harboring an expression vector of *wzx,* p*wzx* | This study |
| Δ*p49* | P49 gene::pKNOCK-Km$^r$ | (2) |
| Δ*hm3359* | *hm3359* (*gspE2*)::pKNOCK-Km$^r$ | (2) |
| Δ*hm3358* | *hm3358* (*gspF2*)::pKNOCK-Km$^r$ | (2) |
| Δ*hm3354* | *hm3354* (*gspK2*)::pKNOCK-Km$^r$ | (2) |
| Δ*hm3349* | *hm3349* (*gspD2*)::pKNOCK-Km$^r$ | (2) |
| Δ*hm3345* | *hm3345* (*gdpD*)::pKNOCK-Km$^r$ | (2) |
| Δ*hm3344* | *hm3344*::pKNOCK-Km$^r$ | (2) |
| Δ*hm3343* | *hm3343* (*wzx*)::pKNOCK-Km$^r$ | (2) |
| Δ*hm3342* | *hm3342* (*nfnB*)::pKNOCK-Km$^r$ | (2) |
| Δ*hm3343*/p*hm3343* | Δ*hm3343* harboring an expression vector of *hm3343,* p*hm3343* | This study |
| ***Escherichia coli*** | | |
| S17-1/λ*pir* | Plasmid donor cell for conjugative transformation of *S. vesiculosa* HM13 and its derivatives | (3) |

**Table S2 Strains and plasmids used in this study (continued)**

| | | |
|---|---|---|
| **Plasmids** | | |
| pKNOCK-Km$^r$ | A gene knockout plasmid, RP4 *oriT* and R6K γ-*ori*; Km$^r$ | (4) |
| pK*pyrF*$^{HM13}$ | A gene knockout plasmid for *pyrF*$^{HM13}$ | This study |
| pKKP | A gene knockout plasmid carrying *pyrF* of *S. livingstonensis* Ac10 | (5) |
| pKKPH | A gene knockout plasmid carrying *pyrF* of *S. vesiculosa* HM13 (*pyrF*$^{HM13}$) | This study |
| pKKPH-*p49* | *pyrF*$^{HM13}$-based gene-deletion plasmid of *p49* | This study |
| pKKPH-*hm3343* | *pyrF*$^{HM13}$-based gene-deletion plasmid of *hm3343* | This study |
| pJRD-Cm$^r$ | A derivative of a broad host-range vector, pJRD215, carrying a chloramphenicol resistance gene | (6) |
| p*hm3343* | A gene complementation plasmid of *hm3343*, pJRD-Cm$^r$ harboring *hm3343* and its predicted promoter region | This study |
