## Supplementary material for "Capsular polysaccharide-mediated protein loading onto extracellular membrane vesicles of a fish intestinal bacterium, *Shewanella vesiculosa* HM13": Table S3

**Table S3 Primers used in this study**

| Primers | Description | Sequence (5´ to 3´) |
|---|---|---|
| ***Construction of pyrF-deletion mutant*** | | |
| *pyrF*$^{HM13}$+1500-fw | Amplification of an upstream flanking | CTAGAACTAGTGGATCCCCCATGCTGGTCAGCCCAGTAAATTTCT |
| *pyrF*$^{HM13}$-del2 | region of *pyrF*$^{HM13}$ | TGGAACTCCAGCGGTAATATGGTGTTAC |
| *pyrF*$^{HM13}$-del1 | Amplification of a downstream | ATATTACCGCTGGAGTTCCATTCTAGCAATGATGCCAGCAAAGTACAC |
| *pyrF*$^{HM13}$+1500-rv | flanking region of *pyrF*$^{HM13}$ | ATATCGAATTCCTGCAGCCCTAGTCGGCATATGGATCATCAGCGC |
| *pyrF*$^{HM13}$-1.7up-fw | Confirmation of the genome insertion | AGTATGTTCCACACTTTAAGCCGGG |
| pKNOCK-0.15dw-rv | of pK*pyrF*$^{HM13}$ | CTACGTGTTCCGCTTCCTTTAGCA |
| ***Construction of pKKPH*** | | |
| *pyrF*$^{HM13}$-fw | Amplification of *pyrF*$^{HM13}$ | TAGCGTGATAACAAATCAAACATGTAGCGATTGGTAAATAGCCTTG |
| *pyrF*$^{HM13}$-rv | | CCGCTGGAGTCCTAATGACTGAGAAATCAATTGTAGTCGCCC |
| pKKP-fw | Amplification of a linear fragment of | TTTGTTATCACGCTAACGGTTAGGTTGAC |
| pKKP-rv | pKKP excluding *pyrF*$^{Ac10}$ | TAGGACTCCAGCGGCAATAGCG |
| ***Construction of P49-gene-deletion mutant*** | | |
| pKKPH-invfwd | Amplification of a linear fragment of | GGGGGATCCACTAGTTCTAGAGCG |
| pKKPH-invrev | pKKPH | GGGCTGCAGGAATTCGATATCAAG |
| *p49*_upfwd | Amplification of an upstream flanking | GAATTCCTGCAGCCCGGGTTAACGCTTGAAAAGCGAAAAAAAAG |
| *p49*_uprev | region of P49-coding gene | ACCATCTCCAATTTCATGAAATTTGGACTTACC |
| *p49*_downfwd | Amplification of a downstream | AAATTGGAGATGGTTCGTTTATAGAGGCTAATTTTTATCGGCTTATGGC |
| *p49*_downrev | flanking region of P49-coding gene | ACTAGTGGATCCCCCAAGGAGAATAGTAAACAGGCGAATAACGC |
| ***Construction of hm3343-deletion mutant*** | | |
| *wzx*_upfwd | Amplification of an upstream flanking | CTAGAACTAGTGGATCCCCCTGGGTCTATTGTATCCATAGAGGGCAAAC |
| *wzx*_uprev | region of *hm3343* | TCAATTGTAAACCTTTGCCTAAAAAACACCT |
| *wzx*_downfwd | Amplification of a downstream | AGGCAAAGGTTTACAATTGACCAATTTTACCTTAAAAAGGCGTGAGAAA |
| *wzx*_downrev | flanking region of *hm3343* | ATATCGAATTCCTGCAGCCCCTAAGATCGCCGACCACAACAATAGTACA |
| ***Confirmation of gene deletion*** | | |
| Check_*pyrF*$^{HM13}$fwd | Confirmation of the deletion of | AGTATGTTCCACACTTTAAGCCGGG |
| Check_*pyrF*$^{HM13}$rev | *pyrF*$^{HM13}$ | GGCCAAGCAGGCTAATGCAG |
| Check_*p49*fwd | Confirmation of the deletion of the P49- | AGCAAGTGTAAGGATGTTTTTGCAGG |
| Check_*p49*rev | coding gene | TAGCAACCACGCCGTAAATACACG |
| Check_*wzx*fwd | Confirmation of the deletion of *hm3343* | TAGGGTCTTGCACTACTGTCAGGTTTTTC |
| Check_*wzx*rev | | CAAGCGCACCGTCTAAATAAATCAAATG |

**Table S3 Primers used in this study (continued)**

| | | |
|---|---|---|
| ***Complementation of hm3343-deletion mutant*** | | |
| pJRD-invfwd | Amplification of a linear fragment of pJRD-Cm$^r$ | CGTAATCCATGGATCAAGAG |
| pJRD-invrev | | TAGTATAGTCTATAGTCCGTGG |
| *wzx*up500fwd | Amplification of *hm3343* and its upstream flanking region | CTCTTGATCCATGGATTACGCAGGGCGGTAATATGGTGAATTTTTTTATT |
| *wzx*rev | | ACGGACTATAGACTATACTATCACTTGCCATTGAAGCACA |
| pJRD_CheckFW1 | Confirmation of insertion of *hm3343* into pJRD-Cm$^r$ | GTGCGCCAACTACCAGCT |
| *wzx*_CheckFw2 | | GTGATATTATCTGCGGTATCGAAATTTATGC |
| pJRD_CheckRV | | GCCTGACTGCGTTAGCAATTTAAC |
