## Supplemental procedures for "Capsular polysaccharide-mediated protein loading onto extracellular membrane vesicles of a fish intestinal bacterium, *Shewanella vesiculosa* HM13"

**Construction of a *pyrF*-based gene knockout plasmid and preparation of Δ*pyrF*^HM13^**

A DNA fragment composed of the upstream and downstream flanking regions (1,590 and 1,447 bp, respectively) of *pyrF*$^{HM13}$ was introduced into the SmaI site of pKNOCK-Km$^{r}$ to obtain pK*pyrF*$^{HM13}$. The upstream and downstream flanking regions of *pyrF*$^{HM13}$ and linearized pKNOCK-Km$^{r}$ were amplified using the following primer sets: *pyrF*$^{HM13}$+1500-fw/*pyrF*$^{HM13}$-del2, *pyrF*$^{HM13}$-del1/*pyrF*$^{HM13}$+1500-rv, and pKKPH-invfwd/pKKPH-invrev, respectively (**Table S3**). DNA fragments of the flanking regions and linearized plasmid were ligated using the NEBuilder HiFi DNA Assembly Cloning System kit (New England Biolabs, Japan, Tokyo, Japan). The resulting plasmid, pK*pyrF*$^{HM13}$, was introduced into *E. coli* S17-1/λ*pir* and then transferred to *S. vesiculosa* HM13-Rif$^{r}$ by conjugation as previously described (1). Transformants grown on LB agar plates containing kanamycin (Km) and rifampicin (Rif) were selected. Single colonies were picked up, and plasmid integration into the genome was confirmed by diagnostic PCR with the primer set *pyrF*$^{HM13}$-1.7up-fw and pKNOCK-0.15dw-rv. The colony harboring the integrated plasmid was cultured in 5 mL of LB media at 18 °C until the $OD_{600}$ reached approximately 1.0, and then the cells were spread on LB plates containing 40 μg/mL uracil, 50 μg/mL Rif, and 1 mg/mL 5-fluoroorotic acid (5-FOA). After two rounds of selection with 5-FOA, the deletion of *pyrF*$^{HM13}$ from the genome was confirmed by diagnostic PCR using the following primer set: Check_*pyrF*$^{HM13}$fwd and Check_*pyrF*$^{HM13}$rev.

**Construction of a gene-deletion plasmid, pKKPH**

A *pyrF*$^{HM13}$-based gene-knockout plasmid, pKKPH, was constructed. The *pyrF* gene of *S. livingstonensis* Ac10 (*pyrF*$^{Ac10}$) in the pKKP plasmid (2) was replaced by the *pyrF*$^{HM13}$ gene (*hm2280*) of *S. vesiculosa* HM13. A 699-bp DNA fragment containing the entire *pyrF*$^{HM13}$ was amplified from the genomic DNA of *S. vesiculosa* HM13 by PCR with a primer set of *pyrF*$^{HM13}$-fw and *pyrF*$^{HM13}$-rv. Then, a 2.1-kbp fragment of pKKP, except for *pyrF*$^{Ac10}$, was amplified using a primer set of pKKP-fw and pKKP-rv. The two PCR products were purified and ligated using a NEBuilder HiFi DNA Assembly Cloning System kit. The recombinant plasmid was introduced into *E. coli* S17-1/λ*pir*. Transformants were selected on LB plates containing 50 μg/mL Km. The construction of pKKPH was confirmed by diagnostic PCR using the primer set *pyrF*$^{HM13}$-fw and *pyrF*$^{HM13}$-rv.
